# Oncogenic H-Ras Reprograms Madin-Darby Canine Kidney (MDCK) Cell-derived Midbody Remnant Proteins Following Epithelial-Mesenchymal Transition

**DOI:** 10.1101/2025.07.17.665448

**Authors:** Adnan Shafiq, Alin Rai, Rong Xu, Maoshan Chen, Wittaya Suwakulsiri, David W. Greening, Richard J. Simpson

## Abstract

Epithelial-mesenchymal transition (EMT) is a highly conserved morphogenic process that allows highly polarized, immotile epithelial cells to transform into motile mesenchymal cells: it is a fundamental cellular process involved in embryonic development, tumour cell metastasis, organ fibrosis and tissue regeneration. To assess the contribution of secreted midbody remnants (MBRs) - a new class of membranous extracellular vesicle (EV) molecularly distinct from exosomes/small EVs – to the EMT process, we conducted a proteomic analysis of MBRs released from Madin-Darby canine kidney (MDCK) cells, and MDCK cells transformed with oncogenic H-Ras (21D1 cells). MBRs were harvested from cell culture media in milligram quantities using a continuous culture bioreactor device and purified using sequential centrifugation and buoyant density gradient centrifugation (OptiPrep™). Gel-MS/MS protein profiling showed MDCK cell-MBRs reflect their epithelial origin (e.g., enriched CDH1, DSP, THBS1, OLCN, EPCAM proteins) and 21D1 cell-MBRs their mesenchymal phenotype (e.g., HRAS, VIM, MMP14, CDH2, WNT5A and enriched invasive and cell motility proteins). Prominent findings were the unique expression of the immune checkpoint protein NT5E/CD73 (ecto-5′-nucleotidase), and ser/thr kinases LIMK1/K2 in 21D1-MBRs (not present in MDCK cell-MBRs), and enrichment in Wnt signalling network proteins. Collectively, our findings suggest MBRs might play a previously unrecognized role in the EMT process.

**Significance:**

1. Epithelial-to-mesenchymal transition (EMT) is a critical cell biological process that occurs during normal embryonic development and cancer progression. Our study describes, for the first time, the large-scale sequential purification of secreted midbody remnants (MBRs) and exosomes/sEVs from the *in vitro* cell line EMT model Madin-Darby canine kidney (MDCK) cells and MDCK cells transformed with oncogenic H-Ras (21D1 cells): GeLC-MS/MS protein profiling identified the repertoire of enriched MDCK-MBR proteins following EMT.
2. MBRs display a proteome profile distinct from sEVs that is enriched with factors of the centralspindlin complex (KIF23.1, KIF4A, INCENP, CEP55, PLK1) and further include components of the mitochondrial network, cytokinesis, microtubule movement, and intercellular connection.
3. In the context of EMT, our data reveal simultaneous activation of EMT signalling pathways in MBRs including signalling receptor binding, regulation of cell differentiation, and Wnt, VEGF and PDGF signalling.
4. We identify several mesenchymal enriched networks in MBRs associated with focal adhesion, cell matrix, kinase activity, and cell shape/organisation, while epithelial derived MBRs are show enriched networks predominately associated with mitochondrial (processing/transport), midbody, and plasma membrane annotation.
5. Our study sheds light on the signalling architecture of MBRs following oncogenic H- Ras-induced EMT: collectively, our data informs ongoing efforts to delineate oncogenic drivers of cancer initiation, progression, and metastasis.

## 1 INTRODUCTION

Epithelial-mesenchymal transition (EMT) is an evolutionary-conserved cellular process that enables highly polarized immotile epithelial cells to lose their epithelial cell attributes such as cell-cell and cell-matrix connectivity and acquire features of mesenchymal cell phenotype such as elongated morphology, increased motility, reduced proliferation rate^1^; this highly- regulated program is critical for both normal physiological and pathological processes such as early embryogenesis, wound healing, tissue regeneration, and cancer progression^2–4^. EMT can be induced in epithelial cancer cells by several stimuli, including activation of proto- oncogenes (e.g., Ras), growth factor receptor pathways (e.g., those driven by transforming growth factor-β (TGFβ), hepatocyte growth factor (HGF) receptor (MET), or epidermal growth factor receptor (EGFR), for example. Multiple cellular regulatory networks have been implicated in triggering EMT such as tyrosine kinase receptors (e.g., epidermal growth factor (EGF), fibroblast growth factor (FGF), connective tissue growth factor, platelet-derived growth factor (PDGF), insulin-like growth factor (IGF) etc.), and signalling pathways involving integrin interactions, Wnt proteins, nuclear factor NK-κβ, and TGF-β ^2,3,5–7^.

It has been shown that soluble protein factors secreted into the extracellular space (the *secretome*) play a critical role in eliciting the EMT programme^3,8,9^. To identify extracellular modulators of the EMT process, which may influence tumour cell state and invasive potential, we previously assessed the secretome (soluble-secreted proteins)^10,11^ and small EVs^12,13^) released from Madin-Darby canine kidney (MDCK) cells, and MDCK cells transformed with oncogenic H-Ras (21D1 cells) – for reviews, see^14,15^ and references therein. These secretome-based proteomic analyses revealed extracellular effectors that coordinate biological responses that enhance cell mobility, reduce cell-cell contact and cell-matrix adhesion proteins, and remodel the extracellular matrix (ECM)^10,16^. Taken together, our findings indicated that H-Ras induced EMT in MDCK cells resulted in extensive reprogramming of the protein repertoire of exosomes/sEVs – revealed by comparing the proteomes of MDCK cell-derived exosomes (MDCK-Exos) and 21D1 cell-derived exosomes (21D1-Exos)^17^ - including the selective remodelling of regulators of metastatic niche formation, and transcription/splicing factors related to EMT ^12^. We further demonstrated that MDCK/21D1 cell derived microvesicles (MVs) - a EV subclass arising from plasma membrane blebbing - has a distinct proteome composition, compared to MDCK/21D1-Exos, and functional invasive capacity following EMT^13^.

To assess the contribution of a newly described EV (secreted midbody remnants, sMBR) to the EMT programme, we conducted a proteomic analysis of MBRs released from MDCK/21D1 cells. MBRs which have a particle diameter range of 200-600 nm are a major class of membranous EVs, released from most cell types, that arise by symmetrical cytokinetic abscission during final phase of mitosis (telophase)^17–20^.

In our present study we conducted large-scale purification of MBRs and small EVs/exosomes released from MDCK/21D1 cells (sEVs), and determined their proteome composition using high sensitivity proteomic profiling. We demonstrate that MBRs released from MDCK/21D1 cells have distinct proteome repertoires compared to their sEV counterparts: they are enriched in cytokinesis factors in midbodies (e.g., KIF23 and RACGAP1^18,21^) and enriched components associated with mitochondrial network and cytokinesis, microtubule movement, and intercellular connection. We further differential abundance of networks associated with key processes of EMT in H-Ras-induced EMT in MDCK cells including signal transduction (HRAS, PLAUR, RGS12), plasticity (TNC, VGF), polarity (MARK1, FRMD4A, MPP1, FZD2), and the EMT process (TEAD1, MCU, ESRP1, ESRP2, PKP3, TJP3). We report EMT-related transcriptional and post-transcriptional regulators, including TEAD1, MCU, ESRP1, ESRP2, PKP3, and TJP3, suggesting broader reprogramming of gene expression and cell adhesion machinery. Findings from this study provide new knowledge on the signalling architecture of MBRs following H-Ras-induced EMT in MDCK cells.

## 2 EXPERIMENTAL PROCEDURES

### 2.1 Continuous cell culture and large-scale purification of MBRs and sEVs/Exos

Madin-Darby canine kidney (MDCK) cells, and MDCK cells transformed with oncogenic H- Ras (21D1 cells) cells^12,13,22^ were cultured as described^12,13,22^. Primarily for large-scale EV production, 3 x 10^6^ MDCK and 21D1 cells were seeded separately in CELLine AD1000 Bioreactor classic flasks (Integra Biosciences) using EV-depleted FCS (centrifuged at 100,000 *g* for 18 hrs) and incubated for two weeks^13,19,21^. EV types were purified from collected culture media^19^. For each collection, MDCK and 21D1 culture media was sequentially centrifuged at 500 *g* for 5 min (to remove floating cells), 2,000 *g* for 10 min (to remove apoptotic body debris) and 10,000 *g* for 30 min at 4 °C as described^19,21^ (see Supplementary Fig. S1). The 10K pellet was subjected to buoyant density (isopycnic iodixanol (OptiPrep™) gradient centrifugation using a top-down manner as described^23^ and buoyant density fraction #9 (1.21 g/mL) was taken for proteomic sample preparation. MDCK/21D1 cell derived sEVs/exosomes were purified from the 100K-10K supernatant, as described^19^, and fraction #7 (OptiPrep™ buoyant density, 1.10 g/mL) was taken for proteome analyses.

### 2.2 Protein quantitation and western blot analysis

Protein quantification of purified EV samples, and western blot analyses were determined as described^18,21,24^. For Western blot analysis membranes were probed with primary antibodies (anti-mouse ALIX, 1:1,000, Cell Signalling, Cat. No. 2171), (anti-mouse TSG101, 1:1,000, BD Bioscience, Cat. No. 612697), (anti-rabbit GAPDH, 1:3,000, Cell Signalling, Cat. No. 2118), (anti-mouse KIF23, 1:1,000, Santa Cruz Biotechnology, Cat. No. sc-390113), (anti- TGM2, 1:1000, Invitrogen, Cat. No. MA5-12739), (anti-beta actin, 1:1000, Invitrogen, Cat. No. MA1-91399), (anti-mouse RACGAP1, 1:1,000, Santa Cruz Biotechnology, Cat. No. sc- 271110) according to manufacturer’s instructions. The secondary antibodies (IRDye800 goat anti-mouse IgG (Cat. No. AP308P) or IRDye700 goat anti-rabbit IgG (Cat. No. SAB4600400)) were diluted (1:15,000) and the fluorescent signals were detected using the Odyssey Infrared Imaging System, v3.0 (Li-COR Biosciences, Nebraska USA).

### 2.3 EV biophysical characterisation

Cryo-transmission electron microscopic analyses on EV samples were performed as described^18,25^. MBR and sEV particle diameters were determined by nanoparticle tracking analysis (NTA; NanoSight NS300, Malvern) as described^25^.

### 2.4 Proteomic sample preparation

MDCK/21D1-MBR/ sEV samples (n=3, 10 µg) were lysed in 1% v/v sodium dodecyl sulphate (SDS), 50 mM triethylammonium bicarbonate (TEAB), pH 8.0, incubated at 95 °C for 5 mins and quantified by microBCA (Thermo Fisher Scientific) as described ^26^. Proteomic sample preparation was performed using single-pot solid-phase-enhanced sample preparation (SP3) ^27^ as described^28,29^.

### 2.5 Liquid Chromatography–Tandem Mass Spectrometry

Peptides were analysed on a Dionex UltiMate NCS-3500RS nanoUHPLC coupled to a Q- Exactive HF-X hybrid quadrupole-Orbitrap mass spectrometer equipped with a nanospray ion source in positive, data-independent acquisition mode as described^27,28^ with minor modifications. Peptides were loaded (Acclaim PepMap100 C18 5 µm beads with 100 Å pore- size, Thermo Fisher Scientific) and separated (1.9-µm particle size C18, 0.075 × 250 mm, Nikkyo Technos Co. Ltd) with a gradient of 2–80% acetonitrile containing 0.1% formic acid over 110 min at 300 nL min-1 at 55°C (in-house enclosed column heater). Full scan MS were performed in the m/z range of 350 to 1100 m/z with a 120,000 resolution, using an automatic gain control (AGC) of 3 x 10^6^, maximum injection time of 50 ms and 1 microscan. MS2 was set to 45,000 resolution, 1e6 AGC target and the first fixed mass set to 120 m/z. Default charge state set to 2 and recorded in centroid mode. Total scan windows (65) were performed with staggered isolation window from 350 to 1100 m/z and applied with 28% normalized collision energy as described ^30^. MS-based proteomics data is deposited to the ProteomeXchange Consortium via the MASSive partner repository and available via MASSive with identifier (MSV000093044).

### 2.6 Data processing and informatic analysis

Peptide identification and quantification were performed as described previously using DIA- NN (v1.8) ^31^ against *Canis lupus familiaris* (dog, UP000002254) reference proteome (59,101 entries, downloaded 12-2022). Spectral libraries were predicted using the deep learning algorithm employed by in DIA-NN with Trypsin/P, allowing up to 1 missed cleavage ^31^. The precursor change range was set to 1-4, and the m/z precursor range was set to 300-1800 for peptides consisting of 7-30 amino acids with N-term methionine excision and cysteine carbamidomethylation enabled as a fixed modification with 0 maximum number of variable modifications. The mass spectra were analysed using default settings with a false discovery rate (FDR) of 1% for precursor identifications and match between runs (MBR) enabled for replicates ^27^. The resulting output files contaminants and reverse identifications were removed and further analysed using Perseus (v2.0.7.0). Perseus was applied for downstream data processing and analysis. Data quality cut-off was applied with minimum 50% protein group quantification in at least one group. Hierarchical clustering was performed in Perseus using Euclidian distance and average linkage clustering. Gene ontology (biological process, specific level 10) and KEGG pathways (organism: hsa, pvalue cutoff: 0.05) were analysed based on highly-enriched proteins from the statistical analysis using cluster Profiler^32^ (v.3.11, https://bioconductor.org/packages/release/bioc/html/clusterProfiler.html) in R. Principle component analysis, dot, box, and ridge plots were visualized using ggplot (v.3.3.2, https://ggplot2.tidyverse.org/) in R. Heatmaps were visualized using pheatmap (v.1.0.12, https://www.rdocumentation.org/packages/pheatmap/versions/1.0.12) in R. Venn diagram was generated using a web-based tool (http://www.interactivenn.net/)^33^. KEGG pathway analysis was visualized using pathview (v.3.1.2, http://bioconductor.org/packages/release/bioc/html/pathview.html) ^34^ in R.

## 3 RESULTS

### 3.1 Midbody remnants are secreted from MDCK cells

Previously, we demonstrated that cultured MDCK cells – epithelial cell phenotype - undergoing oncogenic H-Ras induced EMT (21D1 cells) – mesenchymal cell phenotype – secrete proteins and membrane-encapsulated vesicles important for remodelling the extracellular environment and promoting cell motility and invasion^12,35,36^: prominent were extracellular matrix (ECM) proteins, cell migration factors, growth factors, proteases and extracellular vesicles – chiefly, small EVs/exosomes (MDCK-Exos, 21D1-Exos). MDCK cells are highly polarised and exhibit a cobblestone-like morphology, while 21D1 cells have a fibroblast appearance and display a spindle-shaped mesenchymal cell morphology (Fig. 1A). Differential centrifugation and buoyant density gradient separation (iodixanol, OptiPrep™) was used in tandem to purify MBRs and exosomes from MDCK / 21D1 cell culture^19,21^ – see Supplementary Fig. S1 for purification strategy. Milligram amounts of purified MBRs were obtained from culture media (CM) by growing parental cells in a continuous bioreactor culture device (CELLine™AD-1000)^13^: typically, 260 mL culture medium (CM) was harvested from the bioreactor device over 36 days – 20 mL per day, 13 days for each biological replicate, n=3. For each buoyant density gradient centrifugation (iodixanol, OptiPrep™) 12 1-mL fractions were manually collected (Supplementary Fig. S1) and subjected to immunoblot analysis: anti-MKLP1/KIF23 and anti-RACGAP1 antibodies were used to identify MBRs in the resuspended 10K pellet; and anti-ALEX and anti- TSG101antibodies to detect exosomes in the resuspended 100K-10K fractions (Supplementary Fig. S3A-D). An inspection of Supplementary Figure S3A-D showed that fraction #9 in the resuspended 10K pellet (buoyant density ∼1.12 g/mL) is enriched in MBRs, and fraction #7 in resuspended 100K-10K pellet (buoyant density ∼1.10 g/mL). MBRs derived from both MDCK and 21D1 cell lines displayed round membranous vesicle structure (based on cryo-EM, Fig. 1B) with a particle diameters of 50 to 680 nm and NTA showed particle diameters of 300-315 nm: in comparison, MDCK-/21D1-Exos were much smaller (30-195 nm diameter based on cryo-EM, and NTA, exhibited a much smaller diameter range – MDCK-Exos 181.5 nm and 21D1-Exos 157.6 nm (according to NTA analysis). Collectively, these data reveal that MBRs and exosomes/sEVs are biophysically distinct. The yields of MBRs and exosomes/sEVs from 260 mL CM, based on protein quantitation analyses, were in the range 1300-1500 µg (MDCK-/21D1-MBRs) and 315-436 µg for MDCK-/21D1-Exos.

**Figure 1.**
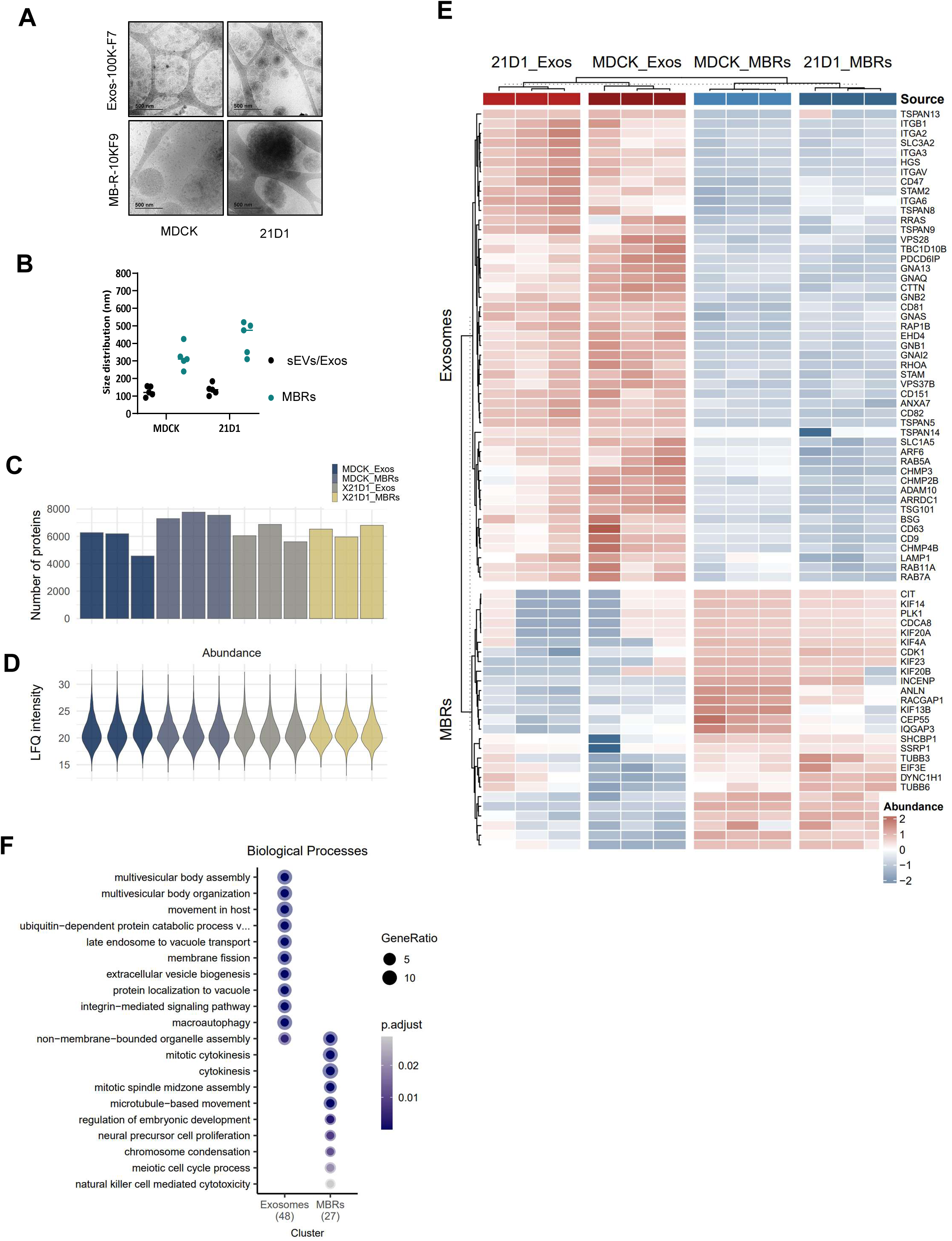
Biophysical and proteome characterisation of MBRs and exosomes oncogenic H-Ras transformed MDCK. A, Cryo-electron microscopy for EV morphology and size distribution. Scale bar, 500 nm. B, Cryo-electron microscopy EV size distribution analysis (each dot represents EV diameter analysis). C, Bar plots for proteins identified and protein distribution based on LFQ normalized intensities for MBRs and exosomes from each cell type (n=3). D, Distribution of protein abundance (LFQ intensity, log2) of each proteome (n=3). E, Hierarchical cluster analysis of MDCK and 21D1 cells and EV subtypes (*P*<0.05, fold change (FC): log2(1.5)), reveals distinct clusters of differential protein abundance based on EV type (MBRs and exosomes). F, Gene Ontology enrichment analysis (ranked, *P*<0.05) of different clusters based on biological process.

### 3.2 Proteome analysis of MDCK- and 21D1-cell derived MBRs and sEVs

Quantitative mass spectrometry using high sensitivity data-independent acquisition ^27^, using stringent peptide/protein identification criteria (1% false discovery rate) identified a total 8737 proteins for all MDCK-/21D1 derived MBRs and sEV preparations analyzed – samples were analyzed in triplicate (n=3). Of these, 4423 proteins were identified in all sample groups in at least n=2 out of 3 biological replicates: protein distribution was based on normalized intensities across all samples - see Figure 1, panels C and D, and Supplementary Table S1). The distinct EV proteomes reported here are highly dynamic, spanning about seven orders of magnitude as measured by their protein abundance (precursor ion intensities), with ACTB, ANXA2, GAPDH, H4C9, and HRAS being top ranked proteins across these datasets based on abundance (intensity) (Supplementary Table S1).

***Hierarchical clustering***. Unsupervised hierarchical clustering of protein expression profiles (Fig. 1E) revealed that MBR and sEV proteomes were distinct: MBR/sEV proteomes were combined for each parental cell type (*P*<0.05, fold change (FC): log2(1.5)). We identified two protein clusters (245 proteins, K-means clustering) with differential abundance between cell origin (MDCK or 21D1) (FC>1.5, p<0.05) in comparison to different EV subtypes (Supplementary Table S2). EV marker distribution was analysed across MBR and exosome populations. sEVs/Exos are enriched in classical markers of small EVs (CD44, CD63, CD9, PDCD6IP (Alix), RAB7A, ITGB1) (Fig. 1E) whereas MBRs were enriched in proteins associated with the centraspindlin complex (KIF23.1, KIF4A, INCENP, CEP55, PLK1) (Fig. 1E). Importantly, these protein markers were associated to EV type, independent of parental epithelial (MDCK) or mesenchymal (21D1) cell origin. Interestingly, the majority of core EV proteins ^37^ including proposed universal markers of exosomes - Syntenin 1 (SDCBP)^37^ and MISEV inclusion protein markers^38,39^ - indicated that EV proteins, including CD47, RAB10, and TSG101 were identified in all EV groups (Supplementary Table S1).

***Gene Ontology (GO)*** GO enrichment analysis revealed enrichment of MBR components associated with cellular location and functions associated with cytokinesis, mitotic spindle midzone assemble, in addition to localisation associated with microtubule network and intercellular bridge (Fig. 1F, Supplementary Fig. S2). In contrast, sEVs were enriched with biological processes associated with multivesicular body assembly, membrane fission, integrin-mediated singling, as well as factors associated with ESCRT complex, focal adhesions, and cell-substrate junctions. Collectively, these data show that MDCK//21D1 cell derived MBRs, and exosomes have distinct proteomes.

### 3.3 Proteins selectively enriched in MBRs released from MDCK cells following oncogenic H-Ras induced EMT

Next, we were curious about whether the MBR proteome from oncogenic H-Ras transformed MBCK cells might reveal new insights into EMT biology. Inspection of the volcano plot in Figure. 2A reveals 267 proteins in 21D1 -derived MBRs are significantly enriched (FC>1.5, p<0.05) and 31/267 are unique to these vesicles (i.e., not seen in MDCK-derived MBRs or sEVs (Fig. 2A, Supplementary Table S1, S3). Of note, networks associated with this 31- protein subset include Interferon Signaling (R-HSA-913531), Cytokine Signaling in Immune system (R-HSA-1280215), and PKR-mediated signaling (R-HSA-9833482), linked with tumor cell regulation, metastasis and EMT process^40–43^. Prominent amongst the 267 proteins enriched in 21D1-MBRs are EMT-related factors WNT5A, STOML3, VGF, MARCKS, and CD70. OIn contrast, ADGRF1, VILL, TSPO, KRT7, and ITGB6 were significantly up- regulated in MDCK-derived MBRs (Supplementary Table S1-S2). We highlight differential abundance of key proteins associated with key processes of EMT in H-Ras-induced EMT in MDCK cells such as signal transduction (up-regulated: HRAS, PLAUR, RGS12), plasticity (up-regulated: TNC, VGF), polarity (up-regulated: MARK1, FRMD4A, MPP1, FZD2), and the EMT process (down-regulated: TEAD1, MCU, ESRP1, ESRP2, PKP3, TJP3) (**Table 1**). Findings from this study provide new knowledge on the signalling architecture of MDCK- MBRs following H-Ras-induced EMT.

**Figure 2.**
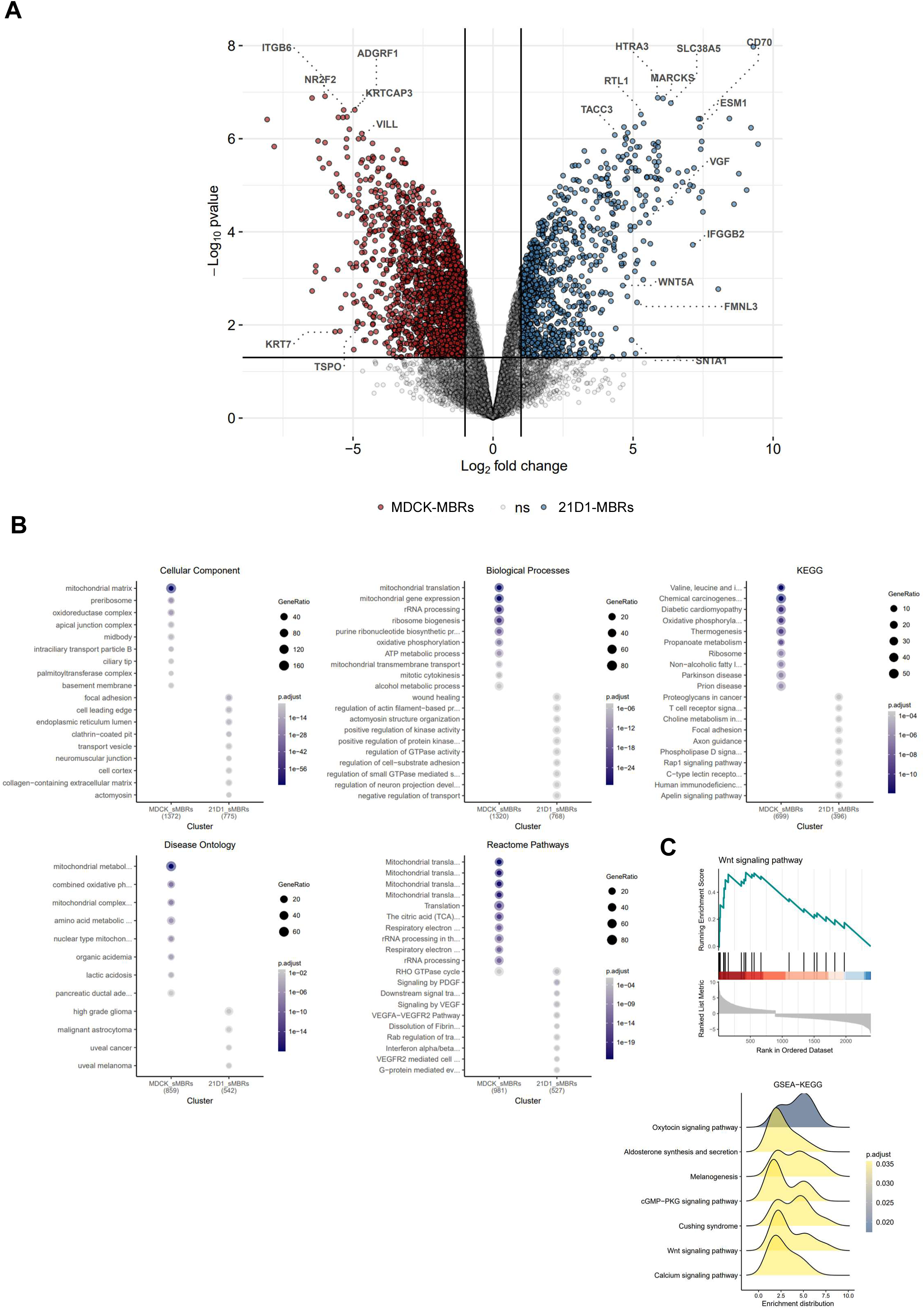
Remodelled proteome cargo of MBRs following H-Ras-induced oncogenic EMT. A, Volcano plot for pair-wise comparison of MBRs proteome in MDCK or 21D1 cells (n=3). Enriched[(blue) or depleted (red), p-value (log10) 1.5 relative to 21D1 MBR. Protein expression non-significant as shown, ns. B, Gene Ontology enrichment analysis (ranked, *P*<0.05) of pairwise comparison of MBRs based on cellular component, biological processes, KEGG pathway analyses, ddisease ontology (DO) in the context of disease, and Reactome analyses. C, Pathway map of Reactome pathways enriched in MBRs (21D1/MDCK, red enrichment in 21D1). Black vertical lines indicate positions where proteins associated with Wnt signalling pathway are in the ranked gene list. Gene Set Enrichment Analysis of KEGG enriched networks in MBRs (p-adj, <0.05), supports enrichment in 21D1-MBRs (yellow) relative to MDCK-MBRs.

Gene Ontology analysis of MDCK-MBRs revealed enrichment of cellular regions and biological processes associated with mitochondrial matrix, ribosome biogenesis, apical junction complex, midbody, basement membrane, rRNA processing, mitotic cytokinesis, and palmitoyl transferase complex (Fig. 2B). In contrast, GO terms enriched in 21D1-MBRs included processes associated predominantly with EMT and cancer progression, including focal adhesion, cell leading edge, collagen-containing extracellular matrix, wound healing, and the positive regulation of kinase and GTPase activity (Fig. 2B, Supplementary Table S4). Further differential analysis of MBR proteome associated with MDCK-MBRs (FC<-1.5, *p*<0.05) displayed upregulation in various processes related to mitochondrial translation, the tricarboxylic acid (TCA) cycle, and rRNA processing (Fig. 2B). In contrast, for upregulated proteins in 21D1-MBRs (FC>1.5, p<0.05) using KEGG and Reactome pathway analysis revealed categories associated with PDGF and VEGF-A/R signalling pathways, Rab GTPase regulation, interferon alpha/beta responses, fibril dissolution, proteoglycan in cancer, and Rap1 signalling pathway (Fig. 2B). WNT signalling pathway components (WNT5A, PLCB1, CCND1, PORCN, FZD2, CHP1, PRKCA, MAPK9, PPP3CC, PPP2R5B, NFAT5, DVL1) were further identified to be significantly enriched relative to the entire MBR proteome from MDCK and 21D1 cells (Fig. 2C). This was further supported by KEGG analysis using Gene Set Enrichment Analysis (GSEA), which further identified WNT signalling pathway as being enriched in MBRs following H-Ras mediated EMT (Fig. 2C). These data highlight the significant remodelling of MBRs proteome occurs following H-Ras induced EMT (network analysis of enriched biological processes, signalling networks associated with 21D1-MBRs, Fig. 3), to support potential signalling capacity of MBRs in the context of EMT.

**Figure 3.**
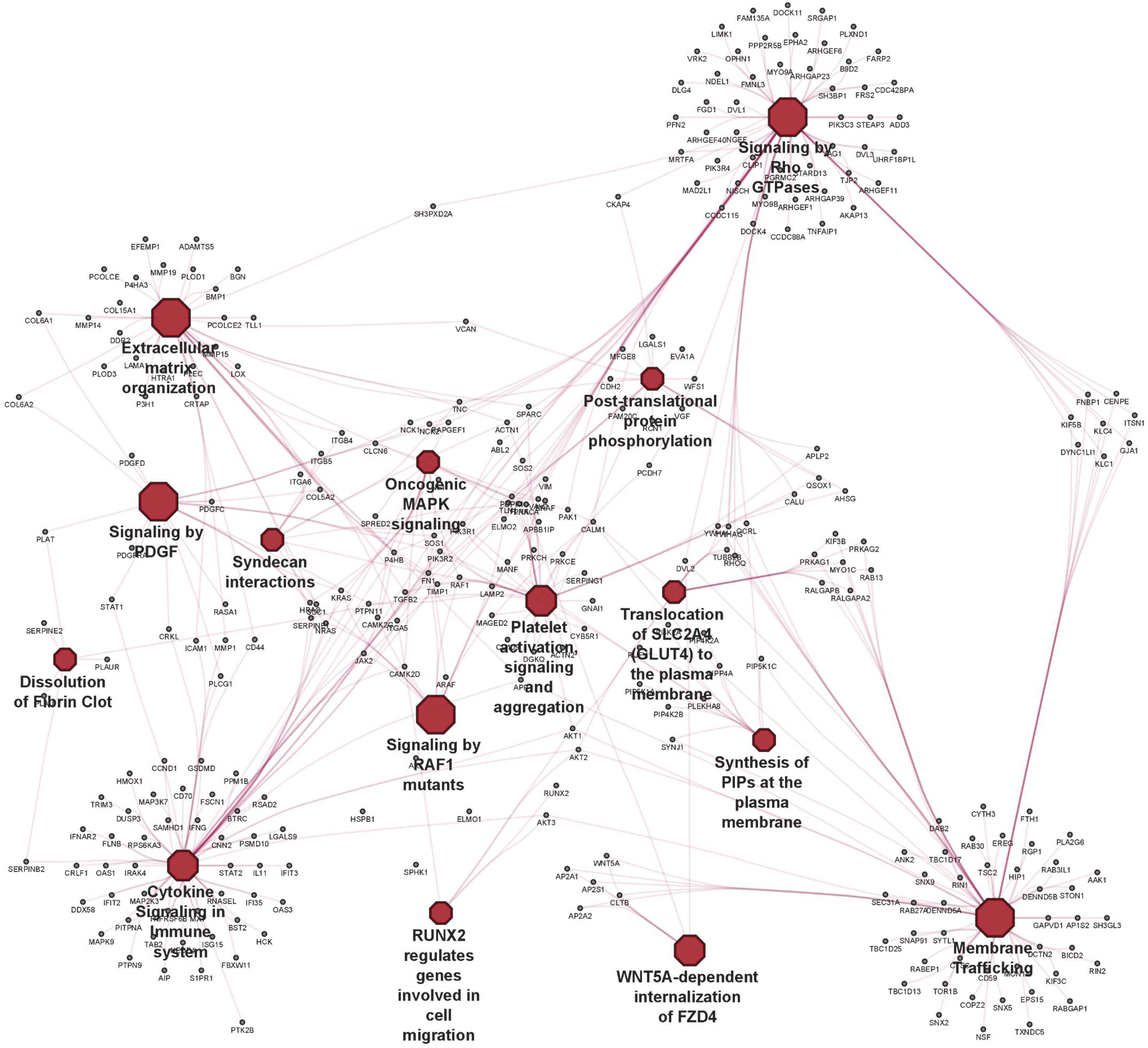
Protein interaction network of 21D1 cell derived-MBRs. STRING protein interaction network for proteins differentially enriched in 21D1-cell derived MBRs based on pair-wise comparison of MBRs proteome relative to MDCK cell derived MBRs. Significantly enriched central terms/pathways/networks are shown in addition proteins associated with each node.

## 4 DISCUSSION

In this study, we describe a large-scale purification procedure for obtaining milligram amounts of highly purified MBRs and exosomes/sEVs from MDCK cells transformed with oncogenic H-Ras. The purification procedure involved continuous culture of MDCK/21D1 cells (CELLine™AD-1000) followed by processing the CM by sequential use of differential centrifugation and buoyant density centrifugation (iodixanol, OptiPrep™). This workflow enabled milligram amounts of EVs for biochemical characterization. High-sensitivity data- independent acquisition (DIA) mass spectrometry was used to obtain protein profiles for MDCK/ 21D1 MBRs and exosomes/sEVs. Our analysis showed that MDCK-MBRs and 21D1-MBRs have distinct protein signatures that distinguish one MBR from another as well as exosomes/sEVs derived from the same parental cell lines. We describe key signaling networks in MBRs for the first time, highlighting oncogenic signaling factors, changes in ECM organization, and associated functions of cytokine signaling, Wnt signaling, protein localization, and membrane trafficking, associated with MBRs following EMT.

Identified proteins common to both MDCK-/ and 21D1 derived from MBRs reflect their respective parental cells: these include classical EMT markers, such as VIM, MMP14, CDH2, WNT5A, and HRAS from 21D1 cells, while proteins associated with MDCK epithelial cells, included CDH1, DSP, OCLN, THBS1, and EPCAM. Proteins enriched in exosomes correlated with known exosomal marker proteins: for example, PDCD6IP (ALIX), CD9/44, and RAB7A, and various tetraspanins and ESCRT related factors such as ESCRT 0 (HGS, STAM1), ESCRT I (MVB12A/B, TSG101, VPS26, VPS37), ESCRT II (VPS25, VPS36) and ESCRT III (CHMP1, CHMP2A/B, CHMP3, CHMP4A/B). Proteins enriched in MDCK-/21D1 derived MBRs included markers related to cytokinesis, such as KIF4A, KIF23, INCENP, CEP55, and PLK1. We also noticed other categories of proteins such as mitochondrial proteins, splicing factors, RNA binding proteins and DNA binding components were significantly enriched in MBRs, consistent with our previous findings for the isogenic colorectal cells lines SW480/SW620 ^18,21^.

Additionally, we identified several integrin components that were significantly enriched in 21D1-MBRs – for example, ITGB1/2/4/5, ITGAV/3/5/6/10. In the case of MDCK-MBRs the only integrins identified were ITGA3 and ITGB6. In addition to their ascribed functions in cancer progression^44,45^, integrins have been shown to be important in exosomal delivery and organotrophic metastasis^46^. Consistent with our finding of integrins in MDCK/21D1- MBRs, Prekeris and colleagues report that secreted midbodies (referred to as ‘midbodysomes’) require integrins and EGF receptor tyrosine kinase signalling for their functionality^18,47^. As expected, GO analysis revealed cytokinesis-related proteins commensurate with MBR biogenesis - such as mitotic spindle midzone assemble, and proteins associated with midbody localisation in the intercellular bridge (e.g., KIF14, KIF4A, KIF23, CDK1 etc).

A salient finding in our study was the remodelling of the MDCK-derived MBR proteome following oncogenic H-Ras induced EMT. Our data analyses support previous findings that EVs comprise signalling factors associated with Wnt and can regulate EMT^12,13,48,49^. Expression of the Wnt family is strongly associated with epithelial differentiation, including capacity to promote EMT and metastasis of pancreatic cancer cells and non-small-cell lung cancer cells^49,50^. Wnt5a is known as a typical non-canonical and Wnt protein is associated with the progress and development in many malignant tumours. Many previous studies have demonstrated that Wnt5a is up regulated in various cancers, including gastric, colorectal, pancreatic, and prostate cancers^51^. WNT5A produced by the TME enables the activation of CDC42, a small GTPase of the Rho family, in hematopoietic stem cells and results in EV release, including sEVs/Exos^52^. However, given that such sEVs were isolated with a direct ultracentrifugation strategy, which may contain other EV subtypes that co-purify (e.g., MBRs) that likely contributed to these Wnt5a findings. Indeed, this is supported by the fact we identify various pro-angiogenic factors including VEGF in both EV types in this study – shown by Ekstrom et al.,^52^ that exosomes/sEVs contain VEGF which is reported to favour angiogenesis, and modulation of adhesion, migration, homing and engraftment.

A major finding in our differential MDCK-/21D1-derived MBR proteome study was the unique identification of NT5E (ecto-5′-nucleotidase, or CD73) and the ser/thr kinases LIMK1 and LIMK2 in 21D1-MBRs (not present in MDCK cell derived MBRs). NT5E is an immune checkpoint protein involved in converting ATP into adenosine, promoting immune escape of tumour cells^53^. Xu and colleagues identified CD73 as highly expressed in gastric cancer linked with poor prognosis, identifying CD73 and adenosinergic signalling within the tumour microenvironment: here, CD73 was also shown to be capable of inducing EMT phenotype during gastric cancer metastasis^54^. Additionally, transcriptome analysis revealed that CD73 modulates the RICS/RhoA pathway extracellularly, leading to LIMK/cofilin phosphorylation inhibition and β-catenin activation^54^. Of note, we identify enriched expression of ser/thr kinases LIMK1 in 21D1-MBRs (relative to MDCK); functionality of this ser/thr kinase as part of the Rho family GTPase signal transduction pathway, involved in microtubule stability and actin polymerisation^55^, and various cell fate processes including cancer development, neurological diseases, and viral infections (reviewed^56^). Importantly, prior studies have identified LIMK1 as a cancer-promoting regulator in multiple organ cancers, such as breast, prostate, and colorectal cancers^57–60^. Our proteome analysis of unique proteins associated with only MBRs from mesenchymal cells (relative to MDCK cells and sEVs) demonstrate involvement and enrichment of proteins associated with immune regulation, including interferon signalling (FANCM, TRIM3, IFIT3, TARBP2), genes involved in invasion (GMPR), and cytokine signalling (STAT5A, TNFRSF6B, FANCM, TRIM3, IFIT3, TARBP2)^3,61^. In the context of EMT and immunoregulation, whether these molecular MBR cargo can transfer their functionalities to other recipient cells warrants further investigation.

In summary, we demonstrate for the first time the large-scale isolation and biophysical/biochemical characterisation of MBRs from MDCK cells following oncogenic H-Ras induced EMT. We highlight distinct proteome signatures between MDCK-/21D1 derived MBRs and exosomes: we show how the MDCK-MBR proteome is extensively reprogrammed following oncogenic H-Ras induced EMT. Our findings demonstrate MBRs as a unique class of EVs – distinct from exosomes - and suggests they may have previously unrecognized roles in cell differentiation, cell polarity and cancer biology.

## AUTHOR CONTRIBUTIONS

AS, and RJS conceived and designed the study. AS, AR, and RX conducted the experiments. AR and DWG performed all proteomic data generation, bioinformatic analyses and interpreted the data. MC and WS provided intellectual input. DWG and RJS supervised the project and, with AS and AR, wrote the manuscript.

## ACKNOWLEDMENTS

AS and RJS acknowledge funding support from La Trobe University. AS was supported by La Trobe University Post Graduate Scholarship. We acknowledge La Trobe University Comprehensive Proteomics Platform for providing infrastructure.

## CONFLICT OF INTEREST STATEMENT

Authors declare no conflicts of interest.

---

**Supp Figure 1.**
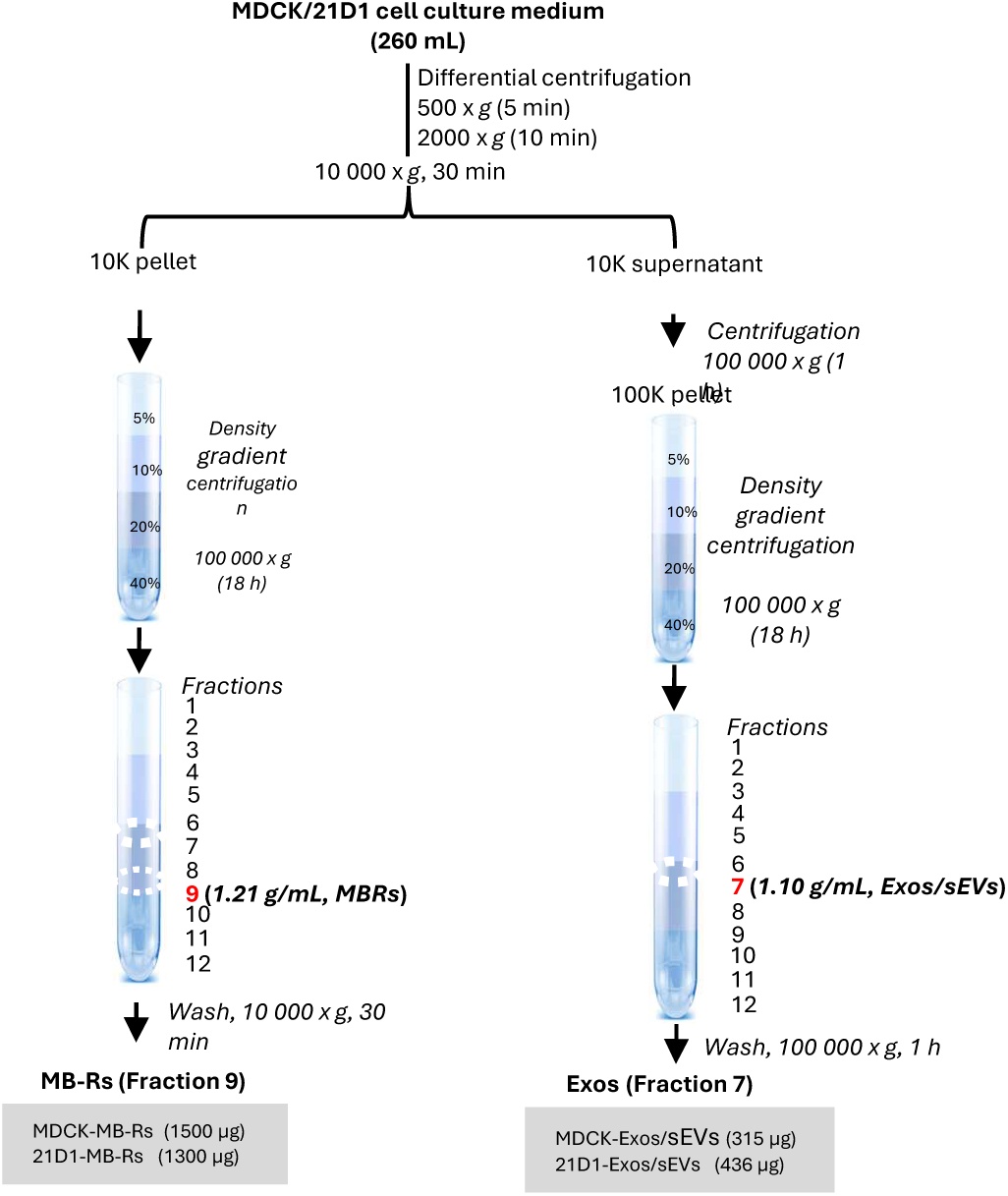

**Supp Figure 2.**
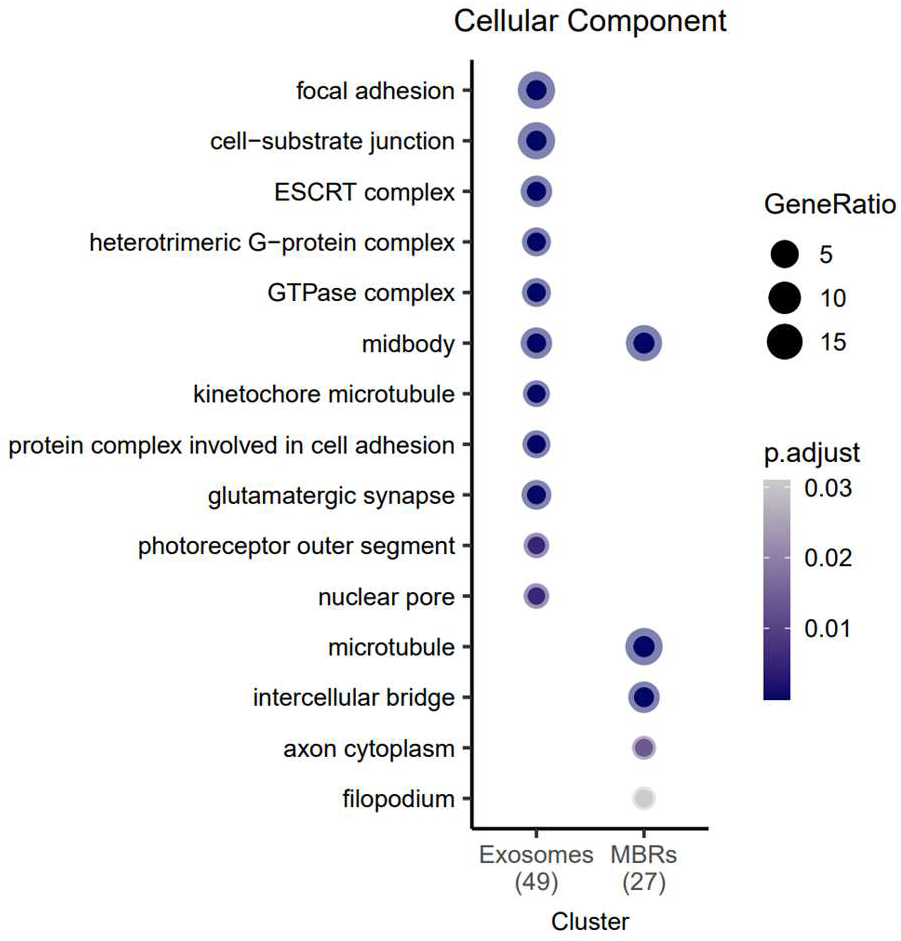

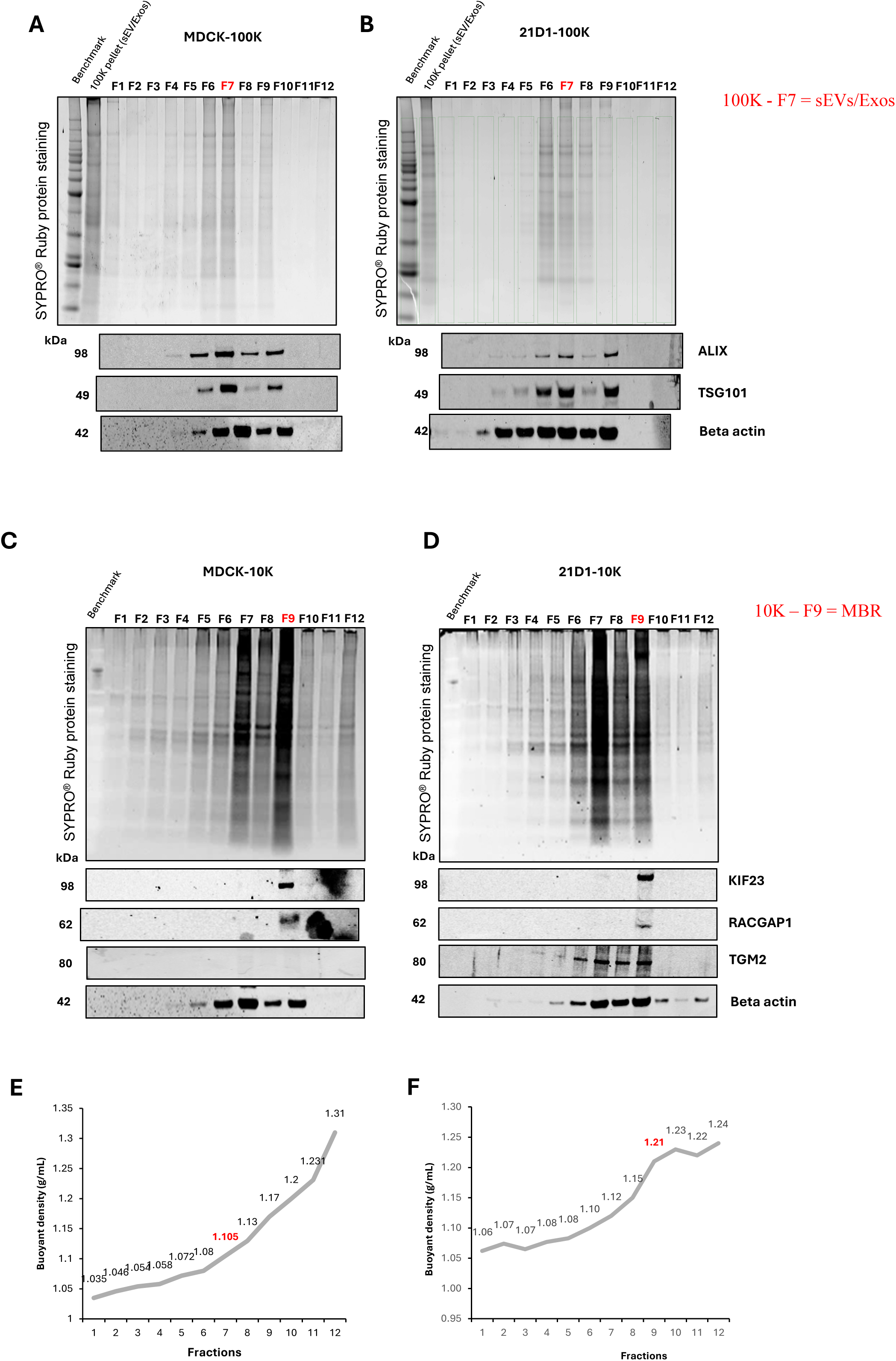

## Notes

### Competing Interest Statement

The authors have declared no competing interest.

